# Divergent myelopoiesis and macrophage polarization underlie host susceptibility to chronic *Chlamydia trachomatis* infection

**DOI:** 10.64898/2025.12.10.693437

**Authors:** Naveen Challagundla, Shivani Yadav, Reena Agrawal Rajput

## Abstract

Chronic genital infection with *Chlamydia trachomatis* is associated with immune suppression and pathogen persistence, but the mechanisms underlying this phenomenon remain poorly defined. Here, we show that chronic *C. trachomatis* genital infection differentially reprograms myelopoiesis and macrophage polarization in genetically distinct mouse strains. In BALB/c mice, infection expanded bone marrow progenitors with enhanced colony-forming potential but skewed their differentiation toward immunosuppressive M_2_ macrophages and myeloid-derived suppressor cells (MDSCs). These cells accumulated in the spleen, blood, and genital tract, secreted IL-10 and TGFβ, and suppressed T cell proliferation, thereby impairing IFNγ-driven clearance. In contrast, C57BL/6 mice exhibited *C/EBPβ*-driven myelopoiesis favoring M_1_ polarization, elevated IL-12 and nitric oxide production, and robust Th1 responses that facilitated bacterial elimination. Together, these findings demonstrate that host genetic background dictates divergent innate programming during *C. trachomatis* infection, and that the balance between M_1_- and M_2_/MDSC-dominated myelopoiesis determines whether infection resolves or progresses to chronicity.

## 1. Introduction

*Chlamydia trachomatis* is the most common bacterial cause of sexually transmitted infections worldwide and frequently infects the female genital tract, where it is often asymptomatic. While most primary infections are resolved by host immune responses, particularly IFNγ-driven clearance, unresolved or recurrent infections can persist and contribute to significant complications, including pelvic inflammatory disease, tubal obstruction, ectopic pregnancy, and infertility(1, 2). Persistent infection is associated with reduced IFNγ responses and the ability of *C. trachomatis* to evade degradative pathways, enabling it to survive within host cells under conditions of chronic inflammation. The mechanisms that underlie immune failure and pathogen persistence in chronic *C. trachomatis* infections are not fully understood, representing a critical gap in our ability to develop effective interventions.

Host genetic background plays a decisive role in shaping susceptibility and resistance to infectious diseases. Inbred mouse strains provide an established framework for dissecting host–pathogen interactions. C57BL/6 mice are widely used for immunological studies due to available genetic tools, whereas BALB/c mice often provide complementary models by exhibiting contrasting immune phenotypes(3). Differences between these strains frequently reflect distinct Th_1_- and Th_2_-associated immune predispositions, which emphasize differential susceptibility across a range of infections, including chlamydial disease(4), Chagas disease(5), helminth infections(6), and leishmaniasis(7). Understanding these genetic variations provide valuable insights into the host-pathogen interactions across diverse infectious diseases. These strain-dependent biases provide a tractable system to investigate how genetic background influences infection outcomes.

In the case of *C. trachomatis*, C57BL/6 mice mount Th_1_-dominated responses that promote bacterial clearance, whereas BALB/c mice often develop chronic infection with regulatory or Th_2_-biased immune profiles (2, 8). Such divergent trajectories point to immune programming as a central determinant of whether infection resolves or persists. However, the mechanisms by which genetic background reprograms innate and adaptive immunity during chronic chlamydial infection remain unclear.

Here we leverage the contrasting immune predispositions of BALB/c and C57BL/6 mice to investigate how genetic background influences systemic immunity during chronic *C. trachomatis* infection, with a particular focus on myelopoiesis and macrophage polarization.

We reveal a novel mechanism whereby host genetic background interfaces with infection-driven soluble mediators to skew myeloid progenitor differentiation into immunosuppressive M_2_ macrophages and myeloid-derived suppressor cells (MDSCs) in BALB/c mice, promoting chronicity. Conversely, C57BL/6 mice exhibit M_1_-biased progenitor programming and inflammatory macrophage responses leading to bacterial clearance. Understanding these pathways provides new mechanistic insight and identifies myeloid lineage manipulation as a promising therapeutic avenue for managing persistent chlamydial infections.

## 2. Results

*Chlamydia trachomatis* induces a relatively weak immune response in murine genital tract infection models, nevertheless, these models remain valuable models for studying mechanisms of chronic infection that mimic human genital infections. As an alternative, infection with *C. muridarum* establishes robust and persistent infection in mice, closely mimicking the upper genital tract pathology observed in women, including hydrosalpinx and infertility. However, significant differences in pathogenicity and host-pathogen interactions among *Chlamydia* species and serovars complicate direct comparisons between the immunopathogenesis of *C. trachomatis* to C. *muridarum*(*9*).

### 2.1 Strain-specific immunity drives divergent outcomes of *C. trachomatis* infection

Previous studies, including our own, have demonstrated that genital C. *trachomatis* infection in mice leads to immunosuppression(2, 10, 11). We have demonstrated that the primary infection is effectively cleared through an IFNγ-mediated inflammatory response involving macrophages and CD4^+^ T cells. In contrast, during chronic infection, IFNγ levels are reduced, coinciding with the appearance of aberrant chlamydial forms and the establishment of persistent or latent infection(2). Our research suggests that differences in susceptibility among strains may contribute to variation in acute versus chronic infections. Notably, C57BL/6 mice generally mount a Th_l_- type of immunity, whereas BALB/c mice preferentially generate a Th_2_-biased response(5–7). Leveraging this knowledge, we have extrapolated chronic infection models in these strains to assess *C. trachomatis* ability to induce systemic immune suppression in the mice. This approach allows for a deeper understanding of how host genetic factors influence immune responses during *C. trachomatis* infections and their implications for chronic disease outcomes. We established chronic infection in BALB/c and C57BL/6 mice by administering repeated inoculations of *C*. *trachomatis* (Fig. 1A). These two strains exhibited varying responses to chronic *C. trachomatis* genital infection. C57BL/6 mice effectively cleared the infection within 15 days, while chronic infection persisted in BALB/c mice until day 20, consistent with previous findings (Fig. 1B). The implications of infection lasting longer than 20 days and their impact on secondary infection are detailed in the study(2). Analysis of systemic cytokine in serum revealed that infection resolution in C57BL/6 mice was associated with elevated levels of inflammatory cytokines: TNFα, IL-1β, and IFNγ (Fig. 1C), whereas BALB/c mice exhibited reduced levels of these cytokines but significantly increased levels of IL-10 and TGFβ compared to infected C57BL/6 mice (Fig. 1C).

**Figure 1:**
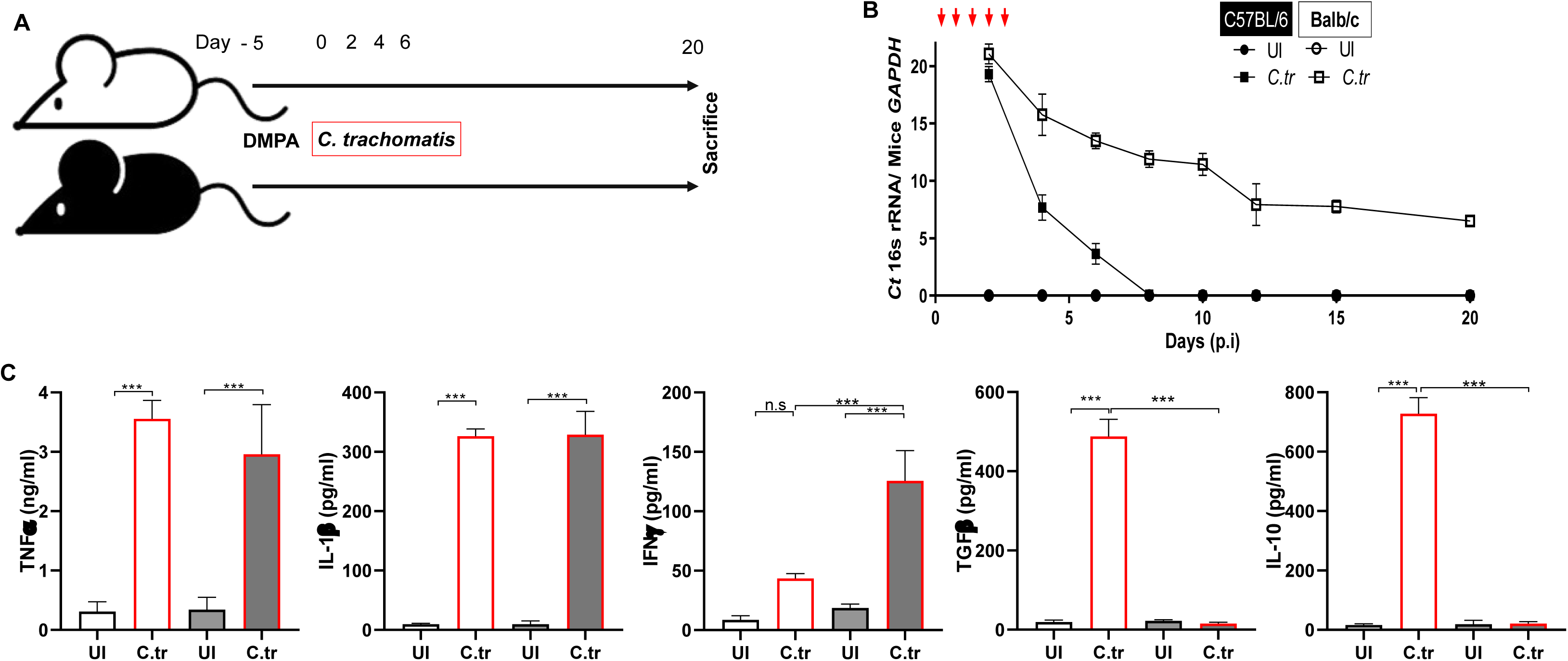
Differential susceptibility of BALB/c and C57BL/6 mice to *C. trachomatis* infection. BALB/c and C57BL/6 mice were injected with DMPA 2.5 mg/kg of body weight (B.W) 5 days prior to infection and inoculated intravaginally with 10⁷ *C. trachomatis* EBs. (**A**) Experimental design. (**B**) Cervical swabs collected every 3 days were analyzed for bacterial burden in McCoy cells. (**C**) Serum cytokine concentrations were measured at day 20 post-infection by ELISA. (n=6) (Mean±SD). one-way ANOVA with Tukey’s multiple comparisons test was performed. ***p ≤ 0.0001; n.s = not significant.

Given that *C. trachomatis* predominantly infects epithelial cells, but macrophages play a pivotal role in orchestrating clearance and immune response at site of infection. To determine the role of macrophages in infection clearance and immune elicitation, we infected the bone marrow-derived macrophages (BMDMs) from BALB/c and C57BL/6 mice separately and compared their responses. BMDMs from C57BL/6 mice displayed a lower bacterial burden compared to BALB/c macrophages (Fig. 2A). This enhanced clearance correlated with robust nitric oxide (NO) production, and elevated TNFα, and IL-12 secretion in macrophages from the C57BL/6 upon *C. trachomatis* infection. In contrast, BALB/c macrophages produced substantially reduced TNFα and IL-12 secretion and lower intracellular NO levels compared to C57BL/6 macrophages (Fig. 2B and C). BALB/c macrophages secreted anti-inflammatory cytokines IL-10 and TGFβ (Fig. 2C). Importantly, the absence of IL-12 production in the BALB/c macrophages during *C. trachomatis*, a cytokine necessary for Th_1_ differentiation, provides mechanistic explanation for the impaired bacterial clearance observed in this strain. Together, these results demonstrate that host genetic background critically shapes the outcome of *C. trachomatis* infection in mice by dictating the balance between pro- and anti-inflammatory immune responses.

**Figure 2:**
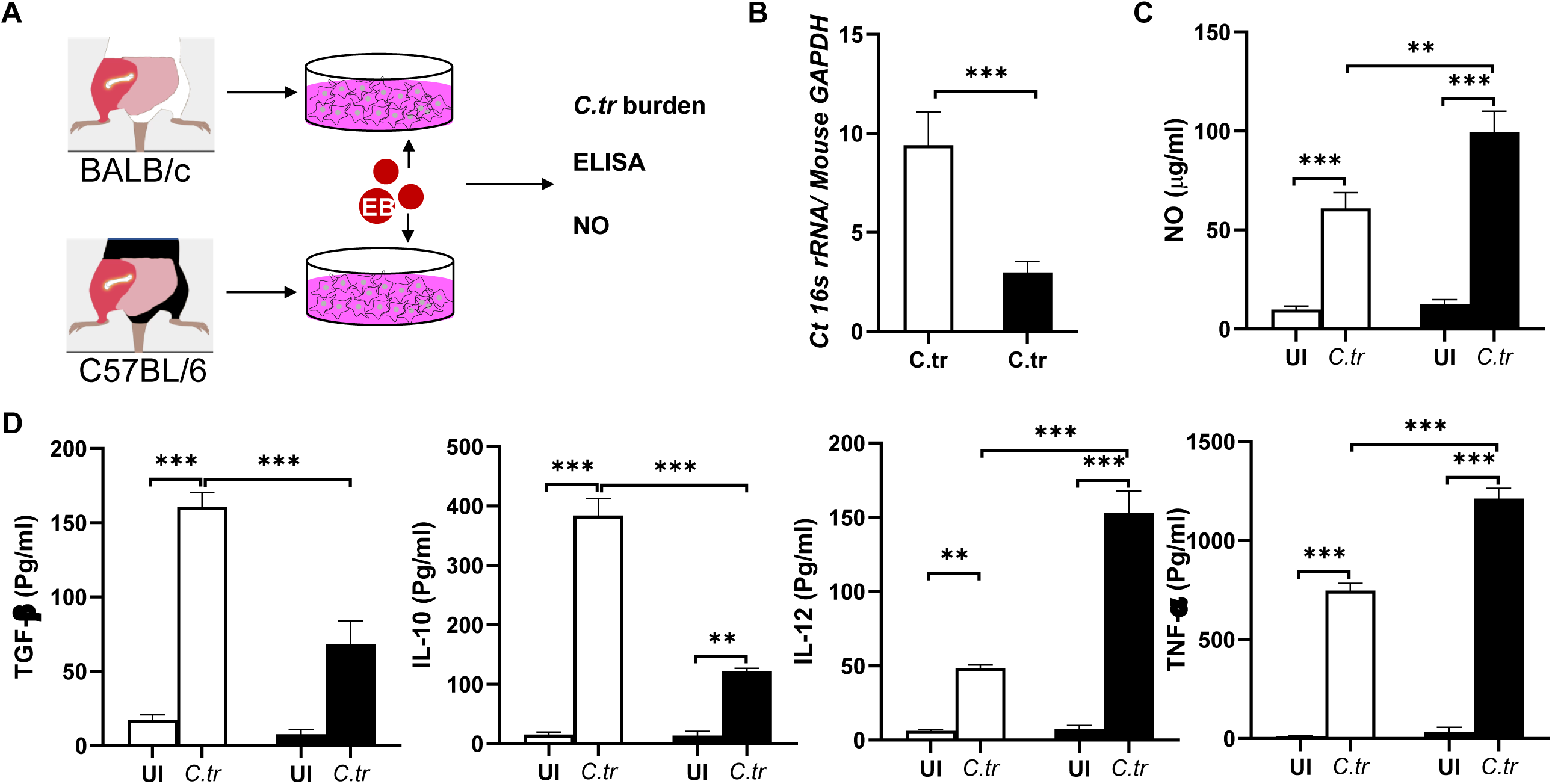
Intrinsic differences in macrophage responses to *C. trachomatis*. Bone marrow–derived macrophages (BMDMs) were generated from BALB/c and C57BL/6 mice using M-CSF and infected with C. trachomatis at MOI 5. (**A**) Experimental design. (**B**) Bacterial burden was quantified by qPCR for C. trachomatis 16S rRNA relative to mouse GAPDH post 24 h post-infection (p.i). (**C**) Nitric oxide levels in supernatants at 12 h p.i were measured using Griess reagent. (**D**) Cytokine secretion at 24 h post-infection was quantified by ELISA. (n=3) (Mean±SD). Student t test was performed in B. one-way ANOVA with Tukey’s multiple comparisons test was performed in C and D. **p ≤ 0.001; ***p ≤ 0.0001; n.s = not significant.

### 2.2 Chronic infection drives distinct macrophage polarization programs in susceptible and resistant strains

Macrophage-derived IL-10 following *C. trachomatis* infection is indicative of polarization toward an M_2_ phenotype. In our previous work, we reported an abundance of M_2_ macrophages at sites of chronic *C. trachomatis* genital tract infection, although their systemic origin was not investigated(2). To address this, in the current study we investigated the macrophage polarization in genital tract tissue, blood and the spleen of BALB/C and C57BL/6 mice during chronic *C. trachomatis* infection. Consistent with our earlier findings, chronic infection in BALB/c mice exhibited a predominance of M_2_ macrophages at the genital tract compared to M_1_ macrophages (Fig 3A). In contrast, macrophages from C57BL/6 mice lacked detectable expression of canonical M_1_ (CD86) or M_2_ (CD206) markers within the female genital tract (Fig 3A and B). However, these macrophages displayed comparatively higher CD40 expression (Fig 3C). The predominance of M_2_ macrophages in the female genital tract of BALB/c mice may be attributed to both local recruitment and systemic generation, as these cells were also abundant in the blood and spleen of infected BALB/c mice (Fig 3D). Indeed, 80% of circulating monocytes in the blood of infected BALB/c mice displayed an M_2_ phenotype compared to 50% in C57BL/6 mice (Fig 3E). Despite this polarization bias, circulating monocytes from both strains showed increased CD40 expression relative to uninfected controls, indicative of infection-induced immune activation (Fig 3F). Splenic macrophage populations further underscored these strain-dependent differences. BALB/c spleens contained a high frequency of M_2_-polarized CD206⁺ macrophages, whereas C57BL/6 spleens harbored an increased proportion of M_1_ macrophages with elevated CD86 expression (Fig 3G and H). Moreover, splenic macrophages from C57BL/6 mice exhibited markedly higher CD40 levels compared to BALB/c counterparts (Fig 3I), consistent with an enhanced ability to provide T cell co-stimulation. This activation phenotype is in line with the increased systemic IFNγ detected in C57BL/6 mice, whereas BALB/c mice demonstrated an immunosuppressive profile favoring infection persistence (Fig. 1A–C). These results suggest that strain-specific differences in macrophage polarization extend beyond the genital tract and reflect systemic myeloid programming.

**Figure 3:**
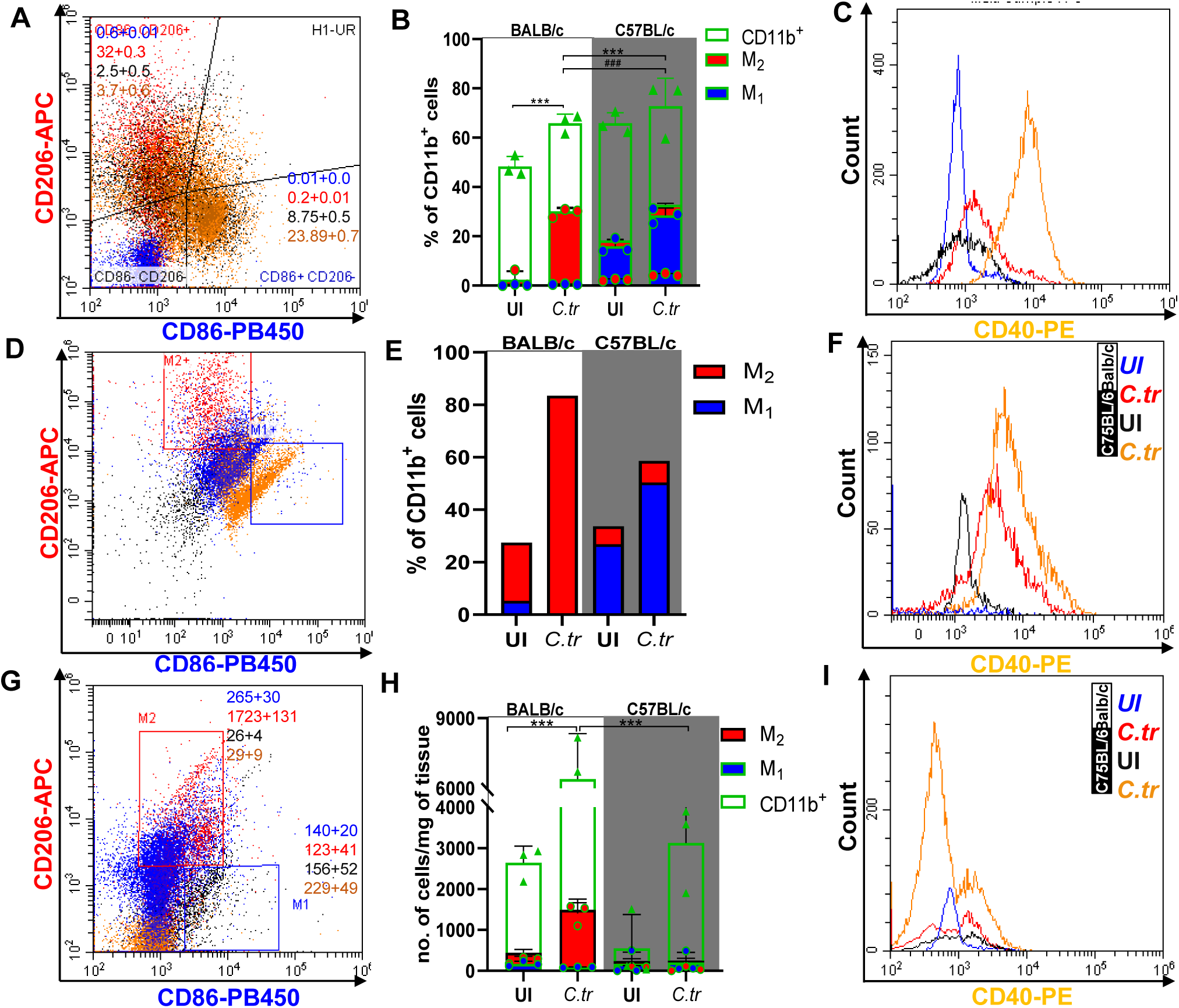
BALB/c mice exhibit increased M_2_ macrophages in systemic and local compartments during infection. RBC-depleted splenocytes from infected and uninfected BALB/c and C57BL/6 mice were stained for macrophage polarization and (**A**) analyzed for M_1_ and M_2_ using flow cytometer and (**B**) percentage of M_1_ and M_2_ macrophages were plotted. (**C**) CD40 expression was analysed on splenic macrophages was analysed using flow cytometry. Blood was collected from control and infected mice. (**D**) Monocytes were collected by histopaque centrifugation, stained, and analyzed for macrophage polarization using flow cytometry and (**E**) percentage of M_1_ and M_2_ macrophages were plotted. (**F**) CD40 expression was analysed on circulating monocytes. Genital tracts from the infected and control mice were digested and (**G**) recruited macrophages were analyzed for macrophage polarization using flow cytometry and (**H**) number of recruited macrophages were plotted. (**I**) CD40 expression was analysed on recruited macrophages. (n=4) (Mean±SD). one-way ANOVA with Tukey’s multiple comparisons test was performed. **p ≤ 0.001; ***p ≤ 0.0001.

### 2.3 Chronic *C. trachomatis* infection induces strain-specific myelopoietic commitment

The spleen acts as a reservoir for myeloid cells, including macrophages, which proliferate and expand in response to infection or injury before migrating to affected tissues. Most circulating and tissue-resident macrophages, however, originate in the bone marrow, where granulocyte–macrophage progenitors (GMPs) and common myeloid progenitors (CMPs) give rise to diverse myeloid populations during infection-driven myelopoiesis(12). Following *C. trachomatis* infection, we observed increased cellularity in the femur, indicative of heightened proliferation of myelopoietic precursors (Fig 4A). This phenomenon was notably more pronounced in BALB/c mice compared to C57BL/6 mice, suggesting a strain-specific differences in the regulation of myelopoiesis during *C. trachomatis* infection. To understand the consequence of chronic infection on myelopoiesis, we isolated bone marrow cells from both BALB/c and C57BL/6 control and infected mice in the presence of M-CSF and IL-3, cytokines known to drive macrophage lineage commitment. Bone marrow cells from infected mice show significantly more colony-forming units compared to the controls, indicating enhanced proliferative capacity and differentiation commitment (Fig 4B). Analysis of these colonies formed in each group revealed varying mRNA levels of transcription factors *C/EBPβ* and *Egr2*, which regulate inflammatory and anti-inflammatory genes in M_1_ and M_2_ macrophages respectively(2, 13). Colonies from the chronic *C. trachomatis*-infected C57BL/6 mice showed elevated levels of *C/EBPβ* mRNA levels consistent with M_1_ macrophage commitment (Fig 4C), whereas colonies from the BALB/c showed upregulated *Egr2* levels suggesting their M_2_-like properties (Fig. 4C). Consistently, analysis of infection stage showed that acute and chronic infections also imposed distinct differentiation programs in the bone marrow, reflected by altered expression of Egr2, C/EBPβ, and CD206 (Supplementary Fig. 1). This transcriptional bias in bone marrow derived cells from different strains was reflected at the functional level as well: BALB/c-derived colonies secreted higher amounts of IL-10 and TGFβ, while C57BL/6 colonies produced elevated IL-12, mirroring the cytokine signatures of M_2_ and M_1_ phenotypes, respectively (Fig 4D). These results are consistent with our previous observation that BALB/c bone marrow-derived macrophages upregulate Egr2 upon *C. trachomatis* infection(2). Together, these results demonstrate that chronic infection drives enhanced myelopoiesis, but bone marrow progenitors are differentially programmed across strains: BALB/c progenitors are skewed toward an immunosuppressive M_2_-like fate, while C57BL/6 progenitors favor an inflammatory, M_1_-like trajectory.

**Figure 4:**
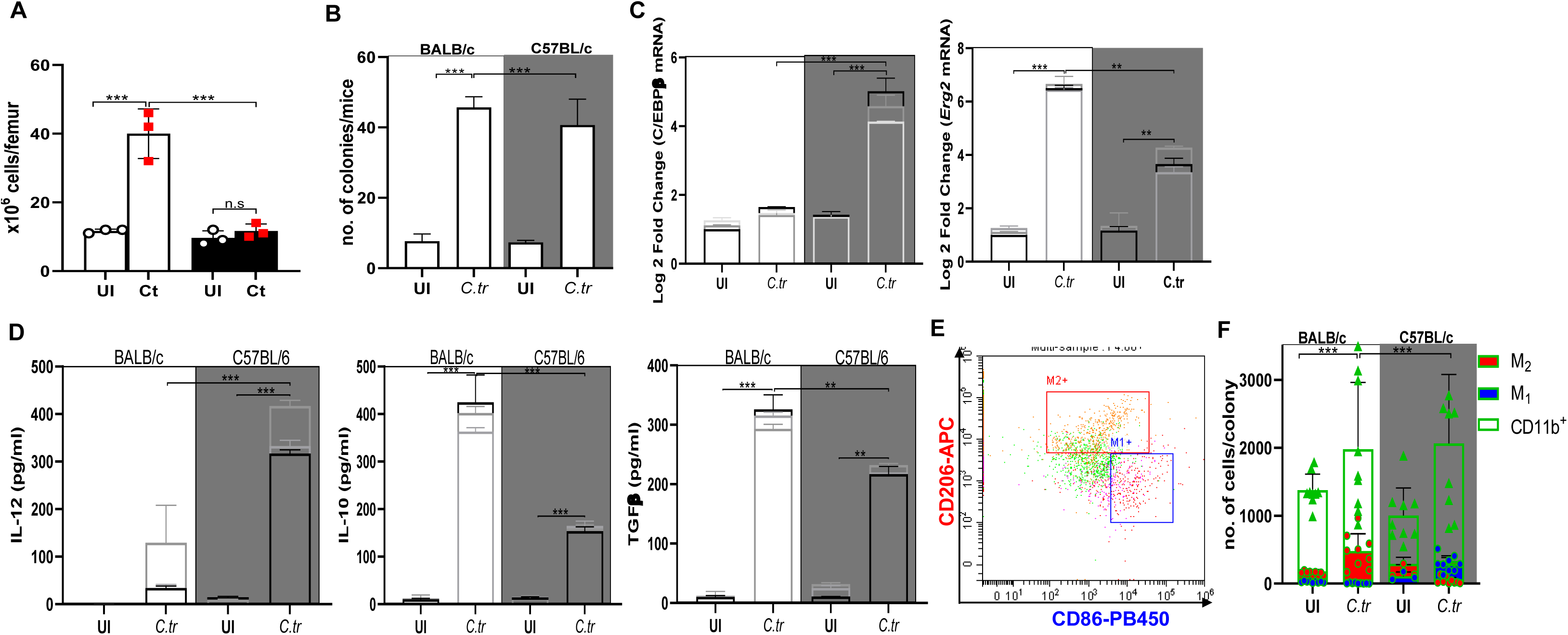
Chronic *C. trachomatis* infection reprograms myelopoiesis in a strain-dependent manner. Bone marrow was harvested from femurs of infected and control BALB/c and C57BL/6 mice. (**A**) Absolute total cell counts per femur. (**B**) Bone marrow cells were cultured in in Iscove’s Modified Dulbecco’s Medium (IMDM) with 0.8% methylcellulose supplemented with IL-3 and M-CSF, and colony-forming units were enumerated at day 6. (**C**) Randomly selected colonies were analyzed for the expression of transcription factors C/EBPβ and Egr2 by qRT-PCR. (n=3). (**D**) Cytokine secretion profiles (IL-10, TGFβ, IL-12) were measured from supernatants of individual colonies after 24 h culture by ELISA Each bar graph represents 3 random colonies produced from a mice. (**E**) Polarization of macrophages within randomly picked colonies was assessed by flow cytometry (n=3). (Mean±SD). one-way ANOVA with Tukey’s multiple comparisons test was performed. **p ≤ 0.001; ***p ≤ 0.0001.

### 2.4 Chronic *C. trachomatis* infection-induced cytokines regulate bone marrow macrophage differentiation

We next examined whether soluble factors released during infection influence bone marrow differentiation, bone marrow cells were exposed to M-CSF in the presence of serum from infected or control animals. Colonies derived from infected BALB/c mice showed a predominance of CD11b⁺ CD206⁺ macrophages, confirming a commitment toward the M_2_ phenotype (Fig 5A and B). In contrast, colonies from infected C57BL/6 mice contained higher numbers of undifferentiated CD11b⁺ macrophages that failed to acquire clear M_1_ or M_2_ polarization markers (CD86, CD206). Although *C. trachomatis* infection generally enhanced macrophage differentiation, serum from infected BALB/c mice was particularly effective in driving M_2_ polarization, suggesting the presence of soluble immunosuppressive mediators. Indeed, BALB/c serum contained elevated levels of TGFβ, IL-1β, and IL-10, along with reduced IFNγ, all of which are known modulators of myelopoiesis (Fig. 1C). In contrast, serum from C57BL/6 mice, which had effectively cleared the infection, exhibited reduced cytokine levels and limited ability to promote macrophage differentiation or skewing, consistent with their restrained myelopoietic response. Together, these findings demonstrate that chronic *C. trachomatis* infection induces systemic myelopoiesis accompanied by strain-specific macrophage lineage polarization. BALB/c bone marrow progenitors display enhanced proliferation and skewing toward an immunosuppressive M_2_ phenotype, driven in part by serum-derived soluble factors, whereas C57BL/6 progenitors favor an M_1_-like program characterized by C/EBPβ induction but limited systemic expansion. These results indicate that soluble mediators released into circulation during chronic infection feed back onto the bone marrow to influence myeloid differentiation, thereby linking peripheral infection with systemic immune programming. While BALB/c serum promotes M_2_-biased differentiation and systemic immunosuppression, C57BL/6 serum exerts limited influence, consistent with efficient infection clearance and restrained myelopoiesis.

**Figure 5:**
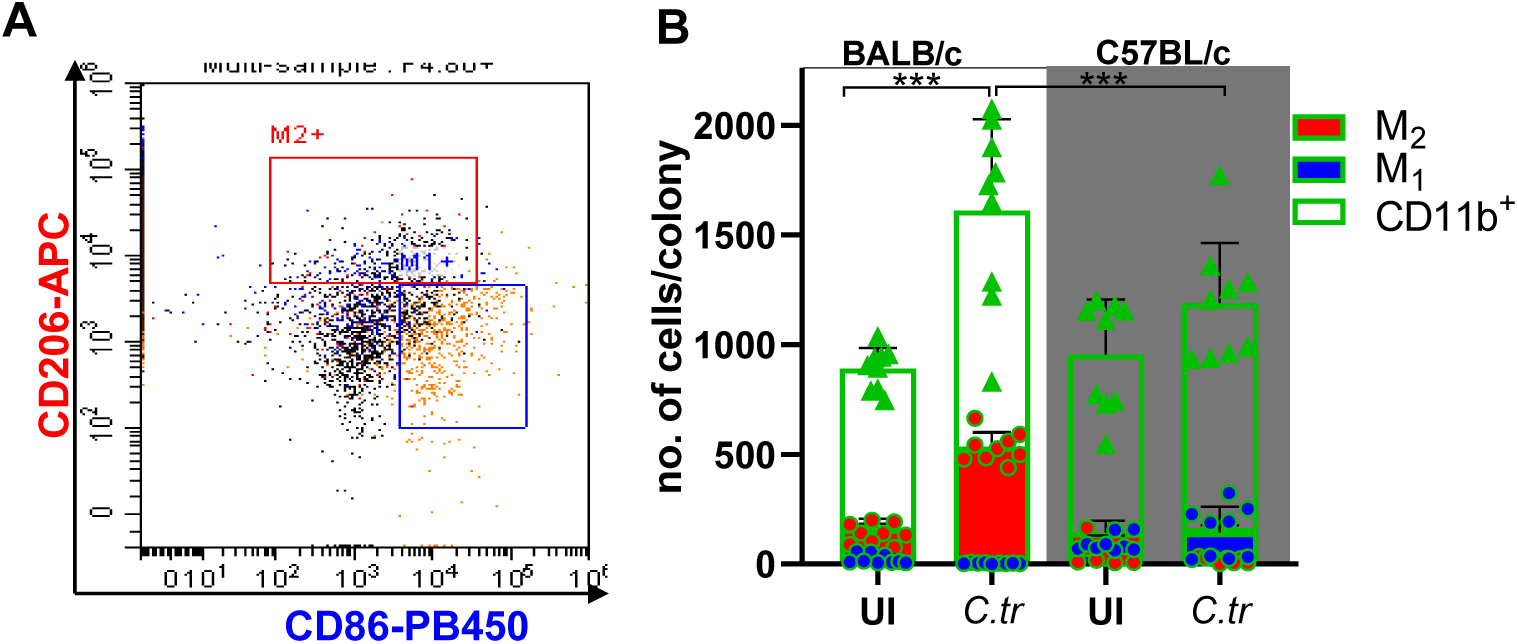
Serum-derived soluble factors from infected mice reprogram bone marrow differentiation. Bone marrow cells from BALB/c and C57BL/6 mice were cultured in IMDM with 0.8% methylcellulose containing IL-3 and M-CSF in the presence of serum obtained from control or infected mice. (**A**) Representative colonies were harvested after 6 days and analyzed by flow cytometry for macrophage polarization using CD11b, CD86, and CD206 markers. (**B**) Frequencies of M_1_- and M_2_-like macrophages from each group were quantified. (Mean±SD) one-way ANOVA with Tukey’s multiple comparisons test was performed. ***p ≤ 0.000.1.

### 2.5 Chronic infection reprograms myelopoiesis to generate immunosuppressive MDSCs

In addition to macrophage differentiation, chronic infection alters myeloid fate by diverting precursors into immature myeloid-derived suppressor cells (MDSCs), a lineage known to exert potent immunosuppressive functions(14). MDSCs arise from both bone marrow and splenic myelopoietic compartments and are broadly classified into monocytic M-MDSCs (CD11b⁺Ly6C⁺) and granulocytic G-MDSCs (CD11b⁺Ly6G⁺). Flow cytometric analysis revealed increased accumulation of MDSCs in the femur of *C. trachomatis*-infected mice (Fig. 5A). BALB/c mice infected with *C. trachomatis* exhibited a marked expansion of M-MDSCs expressing Ly6C and Ly6G in bone marrow, whereas C57BL/6 mice displayed a comparatively modest increase, predominated by G-MDSCs. In the spleen, infection also increased MDSCs, particularly in C57BL/6 mice upon infection with *C. trachomatis*, but the overall abundance remained higher in BALB/c mice. Circulating MDSCs were elevated in both strains, with BALB/c mice showing significantly higher levels than C57BL/6 mice (Fig.5C).

The strain-specific enrichment of MDSCs correlated with differences in macrophage polarization and T cell function. BALB/c mice, characterized by high frequencies of M-MDSCs, also showed increased numbers of M_2_ macrophages in both the spleen and genital tract. In contrast, the comparatively lower abundance of MDSCs in C57BL/6 mice coincided with reduced M_2_ macrophage polarization and enhanced T cell activation, as reflected by increased IFNγ secretion. Together, these findings demonstrate that chronic *C. trachomatis* infection drives strain-dependent expansion of MDSCs, with infection-derived soluble factors not only directing macrophage polarization but also reprogramming myelopoiesis toward MDSC generation, thereby reinforcing systemic immunosuppression.

### 2.6 Divergent T cell responses reflect macrophage polarization states

Innate immune cells, particularly macrophages and dendritic cells, transport antigen to the draining lymph nodes, where they prime T cells and shape downstream effector responses. Activated T cells then secrete effector cytokines critical for controlling infection. Previous studies have established that BALB/c and C57BL/6 mice display distinct immunological predispositions, with BALB/c favoring Th_2_/regulatory responses and C57BL/6 favoring Th_1_ polarization(15–17). To determine whether infection-driven alterations in myelopoiesis and macrophage polarization influence adaptive immunity, we analyzed cytokine secretion from T cells isolated from the inguinal lymph nodes of control and infected mice.

In C57BL/6 mice, infection triggered robust T cell proliferation accompanied by elevated secretion of IL-2 and IFNγ compared to controls or infected BALB/c mice (Fig.6A and B). This enhanced IFNγ response correlated with the presence of M_1_-polarized macrophages in these mice, characterized by elevated IL-12 secretion and high CD40 expression. Consistent with this, serum IFNγ levels were elevated in infected C57BL/6 mice, suggesting that macrophage-driven co-stimulation promotes effective T cell expansion and reinforces Th_1_ immunity, thereby aiding bacterial clearance. In contrast, T cells from infected BALB/c mice failed to secrete IFNγ and instead produced IL-4, IL-10, and TGFβ (Fig.6B), indicative of a Th_2_ bias. These cytokines are known to promote M_2_ macrophage polarization and suppress effector responses, thereby creating a feedback loop that reinforces the immunoregulatory environment observed in BALB/c mice. The strong propensity toward M_2_ polarization, in combination with immunosuppressive mediator release, likely prevents effective T cell activation and contributes to the persistence of *C. trachomatis* in this strain. Together, these findings show that infection-driven macrophage polarization dictates the effector programming of T cells: C57BL/6 macrophages promote T cell expansion and Th1 differentiation, resulting in robust IFNγ production and bacterial clearance, whereas BALB/c macrophages reinforce regulatory polarization, resulting in immunoregulatory cytokines (IL-10 and TGFβ) production, and sustain chronic infection. These results emphasize that the innate immune predisposition of these inbred strains fundamentally shapes adaptive immunity and thereby determines the outcome and progression of *C. trachomatis* infection.

**Figure 6:**
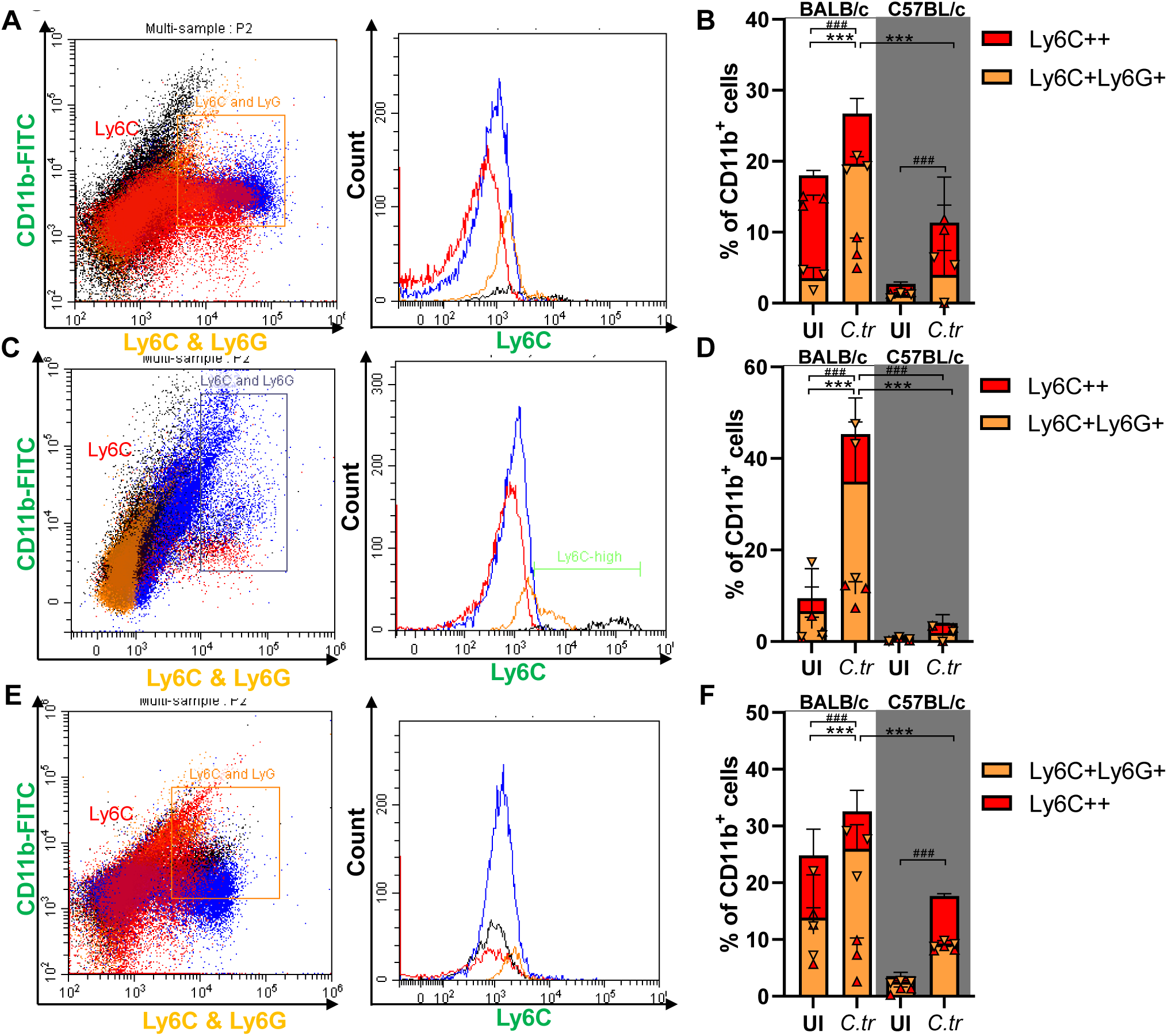
BALB/c mice show greater expansion of MDSCs compared to C57BL/6 mice during infection. Bone marrow cells were obtained from femurs of BALB/c and C57BL/6 mice and cultured in the presence of M-CSF to skew the differentiation. (**A**) Myeloid-derived suppressor cells (MDSCs) were identified by CD11b, Ly6C, and Ly6G expression using flow cytometry CD11b, LY6C and Ly6C\&6G antibodies. (**B**) Percentage of cells were plotted. (**C**) Peripheral blood monocytes were stained for CD11b, Ly6C, and Ly6G to determine circulating MDSCs and percentage of cells were plotted (**D**). (**E**) Splenocytes were analyzed following RBC lysis to quantify MDSC subsets by flow cytometry and percentage of cells were plotted (**F**). (n=4) one-way ANOVA with Tukey’s multiple comparisons test was performed. *** represent p ≤ 0.0001 of Ly6C^+^Ly6G^+^ vs Ly6C^+^Ly6G^+^ and ### represent p ≤ 0.0001 of Ly6C^+^ vs Ly6C^+^.

**Figure 7:**
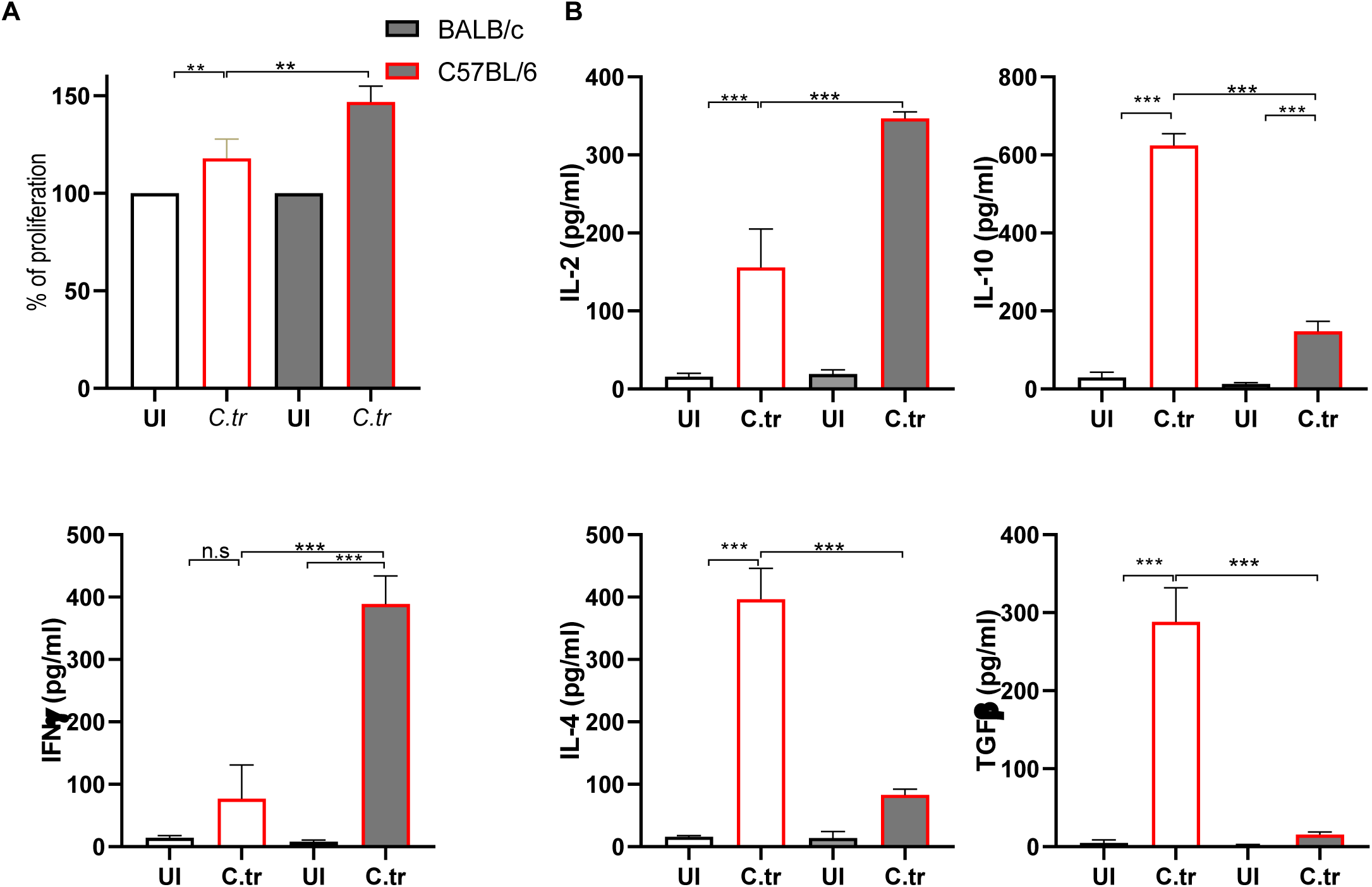
*C. trachomatis* infected BALB/c and C57BL/6 mice differentially show Th_1_ and Th_2_ bias respectively. Inguinal lymph node cells were isolated from infected and uninfected BALB/c and C57BL/6 mice and cultured ex vivo in 96 wells plate. (**A**) T cell proliferation was measured using an MTT assay. (**B**) Cytokine secretion (IL-2, IFNγ, IL-4, IL-10, TGFβ) was quantified in culture supernatants after 48 h stimulation by ELISA (n=4). (Mean±SD). **p ≤ 0.001; ***p ≤ 0.0001; n.s = not significant.

## 3 Discussion

The Th_1_/Th_2_ paradigm has long served as a cornerstone to explain immune responses to various infections, including *Chlamydia trachomatis*. Host genetic background critically influences the balance and thus determines infection outcomes(18). Consistent with earlier studies, BALB/c mice predominantly mount Th_2_/regulatory responses during *C. trachomatis* infection, resulting in pathogen persistence(2, 8), whereas C57BL/6 mice generate a Th_1_-biased response marked by robust IFNγ production and efficient bacterial clearance(8). These contrasting outcomes highlight innate immune predisposition as a fundamental determinant of susceptibility.

Myelopoiesis is dynamically regulated during infection, expanding myeloid precursors to sustain effector responses(12, 18). We show here that chronic *C. trachomatis* infection differentially reprograms myelopoiesis across strains. In BALB/c mice, bone marrow progenitors were skewed toward M_2_ polarization and generated increased myeloid-derived suppressor cells (MDSCs), in part through the action of soluble mediators such as IL-10 and TGFβ. Conversely, progenitors from C57BL/6 mice expressed higher levels of *C/EBPβ* and favored M_1_ differentiation, producing IL-12 and expressing CD40, features that promote inflammatory immunity. These findings extend prior observations that infection-induced cytokines regulate hematopoiesis and highlight the dual contribution of genetic background and soluble mediators in establishing systemic myeloid programming(19–22).

This divergence in macrophage fate was reflected at the functional level. BALB/c macrophages and MDSCs promoted regulatory polarization and impaired effector T cell proliferation, collectively blunting IFNγ responses. In contrast, C57BL/6 macrophages secreted pro-inflammatory cytokines, produced higher nitric oxide, and strongly supported Th_1_ effector functions, consistent with prior reports of strain-specific macrophage programming(23, 24). Accordingly, T cells from C57BL/6 mice proliferated robustly and secreted IFNγ and IL-2, whereas BALB/c T cells secreted IL-4, IL-10, and TGFβ with limited proliferation. Together, these results reveal that macrophage polarization directly imprints T cell programming and thereby shapes infection trajectory.

This study highlights the critical role of myelopoiesis and macrophage polarization in shaping adaptive immunity and determining infection outcomes in *C. trachomatis* infections. We demonstrate that in susceptible BALB/c mice, genetic and cytokine-driven biases favor the expansion of immunosuppressive M_2_ macrophages and myeloid-derived suppressor cells (MDSCs), promoting immune suppression and chronic infection. In contrast, resistant C57BL/6 mice exhibit a robust M_1_-driven inflammatory response that supports Th_1_ immunity and effective bacterial clearance. These findings reveal how host genetic predisposition and infection-induced soluble factors synergize to orchestrate innate-adaptive immune crosstalk, providing mechanistic insight into the dichotomy between bacterial persistence and clearance. Collectively, our work establishes a novel paradigm in which innate immune cell fate decisions, governed by host genetics and systemic cues, dictate divergent disease trajectories in *C. trachomatis* infection. This framework spotlights the myeloid compartment as a pivotal regulator of immune homeostasis during infection and suggests promising therapeutic opportunities to counteract chronic bacterial infection through targeted modulation of myeloid cell development and function.

## 4. Material and Methods

### 4.1. Materials

Medroxyprogesterone 17-acteate and methyl cellulose were procured from Sigma-Aldrich (St. Louis, Missouri, USA). Fluorophore-conjugated antibodies Anti-CD11b FITC (#561688, BD Pharmingen, San Diego, CA, USA), Anti-CD40 PE (#553791, BD Pharmingen), anti-F4/80 APC-R700 (#565787, BD Horizon), Anti-CD206 APC (#17-2061-82, eBioscience, ThermoFischerScientific), Anti-CD68 Pacific blue (#137028, BioLegend, San Diego, CA, USA) Anti-Ly6G&Ly6C (#108412, BioLegend), Anti-Ly6C APC (#565787, BD Pharmingen) were used for flowcytometry. Recombinant mouse (r) IL-10 (#550070), rIL-3 (#554579), rGM-CSF (#554586), rIFNγ (#554587) and rIL-4 (#550067) were procured from BD Pharmingen, San Diego, CA, USA and used for culture assays. rM-CSF (#PMC2044) was obtained from eBioscience, ThermoFischerScientific (San Diego, CA USA). Unless otherwise noted, all additional reagents were purchased from Sigma-Aldrich (St. Louis, Missouri, USA).

### 4.2. Cell culture

Mc Coy cells were procured from the National Center for Cell Sciences, Pune, India and were maintained in McCoy 5A media supplemented with 10% fetal bovine serum (FBS) at 37°C with 5% CO_2_. Bone marrow cells were flushed from femurs of mice under aseptic conditions and cultured in RPMI-1640 medium (Gibco, Thermo Fischer Scientific, USA). To generate bone marrow-derived macrophages (BMDMs), cells were differentiated for 6 days in the presence of rM-CSF, and adherent cells were harvested. Macrophage identity was confirmed by CD11b and F4/80 expression using flowcytometry (CytoFLEX, Beckman Coulter, Brea, CA, USA). Splenic CD4⁺ T cells were isolated from naïve mice following RBC lysis and enrichment with a CD4⁺ T cell isolation kit (BD Biosciences, USA). Purity was verified by CD4 staining and flow cytometric analysis.

### 4.3. *C. trachomatis* propagation and infection

*C. trachomatis* was procured from ATCC, USA and was propagated in McCoy cells, harvested and purified into Sucrose-Phosphate-Glutamine ( SPG) buffer, and stored at - 80°C till further use. Infection stocks were enumerated by inclusion counts. BMDMs were infected at a multiplicity of infection (MOI) 5 in antibiotic-free medium and centrifuged at 750 × g for 30 min to synchronize infection.

### 4.4. Chronic genital *C. trachomatis* infection model

Age-matched female BALB/c and C57BL/6 mice (6-8 weeks old) were maintained under specific pathogen free conditions. All procedures were approved by the Institutional Animal Ethics Committee. To synchronize the estrous cycle, mice received 2.5 mg of medroxyprogesterone acetate (DMPA) subcutaneously 5 days before first infection. Mice were then intravaginally inoculated with 10⁶ inclusion-forming units (IFUs) of *C. trachomatis* in 20ul of SPG buffer(*2*). Infections were repeated every 3 days for a total of five exposures. Control mice were given SPG buffer only. Each experimental group included six mice. Bacterial shedding was monitored by cervical swabs, which were vortexed in HBSS and used to infect McCoy cells. Infection burden was quantified by inclusion counts.

### 4.5. Colony-forming assays for myelopoiesis

Bone marrow cells were collected from the femur resuspended as single-cell suspensions and plated in Iscove’s modified Dulbecco medium (IMDM) supplemented with 0.8% methyl cellulose, 10% FBS, and supplemented with IL-3 (10ng/ml) and M-CSF (20ng/ml). Colonies were enumerated after 6 days(25). Individual colonies were isolated for downstream assays: (i) analysis of cytokine secretion after 24 h culture, (ii) flow cytometric assessment of macrophage polarization, and (iii) gene expression analysis by qRT-PCR. For serum-driven assays, infected- or control-mouse serum (heat-inactivated and 0.2 µm-filtered) was used in place of cytokine supplements to assess the ability of systemic soluble factors to influence colony differentiation.

### 4.6. Flow cytometry

Bone marrow cells, RBC-depleted splenocytes, and peripheral blood monocytes were resuspended in PBS with 2%BSA. Fc receptors were blocked with anti-CD16/32. Cells were stained with CD11b, F4/80, CD206, CD86, or CD68 to evaluate macrophage polarization, or with CD11b, Ly6C, and Ly6G to identify MDSC subsets. After incubation at 4 °C, cells were washed and analyzed using a CytoFLEX flow cytometer (Beckman Coulter). Frequencies of positive cells were calculated per individual mouse(2).

### 4.7. ELISA

Supernatants from macrophage cultures, T cell–macrophage co-cultures, or individual myelopoietic colonies were harvested after 24–48 h and assayed for cytokines (IL-10, IL-12, TGFβ, IFNγ, TNFα) using commercial sandwich ELISA kits(26).

### 4.8. T cell proliferation

Purified CD4⁺ T cells were co-cultured with infected or control macrophages in U-bottom 96-well plates for 48 h Proliferation was assessed using the MTT assay, followed by solubilization with DMSO and absorbance reading at 595 nm. Unstimulated T cells served as negative controls, while CD3/CD28-activated T cells served as positive controls(27).

### 4.9. Quantitative real time PCR

Colonies obtained from the bone marrow were washed in PBS and collected in TRizol (ThermoFischerScientific, San Diego, CA USA). RNA was extracted as described previously(28). Gene expression of C/EBPβ and Egr2 was measured using SYBR Green-based qRT-PCR in StepOnePlus PCR cycler (AB Systems, ThermoFisherScientific, USA). GAPDH served as a normalization control, and expression was represented as fold change relative to controls.

### 4.10. Statistical analysis

All in vivo experiments were repeated twice with at least six animals per group. In vitro experiments were performed in triplicate. All the data are presented as mean ± SD. For comparisons between two groups we used an unpaired two-tailed Student’s t-test; for multiple group comparisons we used one-way ANOVA followed by Tukey’s post hoc test using GraphPad Prism v9.0. *P* values ≤ 0.05 (**) were considered significant.

## 5. Authorship contribution statement

**Naveen Challagundla:** Methodology, software, formal analysis, writing original manuscript. **Shivani Yadav:** Experimentation. **Reena Agrawal Rajput:** Conceptualization, fund management, editing and reviewing the original manuscript and supervision.

## 6. Data availability

All data supporting the findings of this study are available within the article and its supplementary information.

## 8. Acknowledgements

We thank Dr. Manish Nivsarkar for his support in the animal experiments; H. Vora and P. Dalai for their assistance in the laboratory with routine experiments.

## 9. Funding

Naveen Challagundla (67/08/2018-IMM/BMS) and Shivani Yadav’s (45/5/2019-IMM/BMS) fellowship was supported by ICMR, India. Project equipment was partly funded by Fund for Improvement of S&T Infrastructure (FIST), DST India (SR/FST/LS-I/2019/496).

## 10. Ethics declarations

All animal experiments were approved by the Institutional Animal Ethics Committee (IAEC) and were conducted in accordance with institutional and national guidelines for laboratory animal care. The authors declare no competing interests.

## References

1. Lijek RS, Helble JD, Olive AJ, Seiger KW, Starnbach MN. 2018. Pathology after Chlamydia trachomatis infection is driven by nonprotective immune cells that are distinct from protective populations. Proc Natl Acad Sci U S A 115:2216–2221.

2. Challagundla N, Shah D, Dalai SK, Agrawal-Rajput R. 2024. IFNgamma insufficiency during mouse intra-vaginal Chlamydia trachomatis infection exacerbates alternative activation in macrophages with compromised CD40 functions. Int Immunopharmacol 131:111821.

3. Simon MM, Greenaway S, White JK, Fuchs H, Gailus-Durner V, Wells S, Sorg T, Wong K, Bedu E, Cartwright EJ, Dacquin R, Djebali S, Estabel J, Graw J, Ingham NJ, Jackson IJ, Lengeling A, Mandillo S, Marvel J, Meziane H, Preitner F, Puk O, Roux M, Adams DJ, Atkins S, Ayadi A, Becker L, Blake A, Brooker D, Cater H, Champy MF, Combe R, Danecek P, di Fenza A, Gates H, Gerdin AK, Golini E, Hancock JM, Hans W, Holter SM, Hough T, Jurdic P, Keane TM, Morgan H, Muller W, Neff F, Nicholson G, Pasche B, Roberson LA, Rozman J, et al. 2013. A comparative phenotypic and genomic analysis of C57BL/6J and C57BL/6N mouse strains. Genome Biol 14:R82.

4. Darville T, Andrews CW, Jr., Laffoon KK, Shymasani W, Kishen LR, Rank RG. 1997. Mouse strain-dependent variation in the course and outcome of chlamydial genital tract infection is associated with differences in host response. Infect Immun 65:3065–73.

5. Ferreira BL, Ferreira ER, de Brito MV, Salu BR, Oliva MLV, Mortara RA, Orikaza CM. 2018. BALB/c and C57BL/6 Mice Cytokine Responses to Trypanosoma cruzi Infection Are Independent of Parasite Strain Infectivity. Front Microbiol 9:553.

6. Hartmann W, Blankenhaus B, Brunn ML, Meiners J, Breloer M. 2021. Elucidating different pattern of immunoregulation in BALB/c and C57BL/6 mice and their F1 progeny. Sci Rep 11:1536.

7. Barbi J, Brombacher F, Satoskar AR. 2008. T cells from Leishmania major-susceptible BALB/c mice have a defect in efficiently up-regulating CXCR3 upon activation. J Immunol 181:4613–20.

8. Saito S, Nakashima A, Shima T, Ito M. 2010. Th1/Th2/Th17 and regulatory T-cell paradigm in pregnancy. Am J Reprod Immunol 63:601–10.

9. De Clercq E, Kalmar I, Vanrompay D. 2013. Animal models for studying female genital tract infection with Chlamydia trachomatis. Infect Immun 81:3060–7.

10. Marks E, Tam MA, Lycke NY. 2010. The female lower genital tract is a privileged compartment with IL-10 producing dendritic cells and poor Th1 immunity following Chlamydia trachomatis infection. PLoS Pathog 6:e1001179.

11. Batteiger BE, Xu F, Johnson RE, Rekart ML. 2010. Protective immunity to Chlamydia trachomatis genital infection: evidence from human studies. J Infect Dis 201 Suppl 2:S178–89.

12. Challagundla N, Shah D, Yadav S, Agrawal-Rajput R. 2022. Saga of monokines in shaping tumour-immune microenvironment: Origin to execution. Cytokine 157:155948.

13. Veremeyko T, Yung AWY, Anthony DC, Strekalova T, Ponomarev ED. 2018. Early Growth Response Gene-2 Is Essential for M1 and M2 Macrophage Activation and Plasticity by Modulation of the Transcription Factor CEBPbeta. Front Immunol 9:2515.

14. Gabrilovich DI, Nagaraj S. 2009. Myeloid-derived suppressor cells as regulators of the immune system. Nat Rev Immunol 9:162–74.

15. Watanabe H, Numata K, Ito T, Takagi K, Matsukawa A. 2004. Innate immune response in Th1- and Th2-dominant mouse strains. Shock 22:460–6.

16. Garcia-Pelayo MC, Bachy VS, Kaveh DA, Hogarth PJ. 2015. BALB/c mice display more enhanced BCG vaccine induced Th1 and Th17 response than C57BL/6 mice but have equivalent protection. Tuberculosis (Edinb) 95:48–53.

17. Fukushima A, Yamaguchi T, Ishida W, Fukata K, Taniguchi T, Liu FT, Ueno H. 2006. Genetic background determines susceptibility to experimental immune-mediated blepharoconjunctivitis: comparison of Balb/c and C57BL/6 mice. Exp Eye Res 82:210–8.

18. Schultze JL, Mass E, Schlitzer A. 2019. Emerging Principles in Myelopoiesis at Homeostasis and during Infection and Inflammation. Immunity 50:288–301.

19. Christ A, Gunther P, Lauterbach MAR, Duewell P, Biswas D, Pelka K, Scholz CJ, Oosting M, Haendler K, Bassler K, Klee K, Schulte-Schrepping J, Ulas T, Moorlag S, Kumar V, Park MH, Joosten LAB, Groh LA, Riksen NP, Espevik T, Schlitzer A, Li Y, Fitzgerald ML, Netea MG, Schultze JL, Latz E. 2018. Western Diet Triggers NLRP3-Dependent Innate Immune Reprogramming. Cell 172:162–175 e14.

20. Mitroulis I, Ruppova K, Wang B, Chen LS, Grzybek M, Grinenko T, Eugster A, Troullinaki M, Palladini A, Kourtzelis I, Chatzigeorgiou A, Schlitzer A, Beyer M, Joosten LAB, Isermann B, Lesche M, Petzold A, Simons K, Henry I, Dahl A, Schultze JL, Wielockx B, Zamboni N, Mirtschink P, Coskun U, Hajishengallis G, Netea MG, Chavakis T. 2018. Modulation of Myelopoiesis Progenitors Is an Integral Component of Trained Immunity. Cell 172:147–161 e12.

21. Nagareddy PR, Kraakman M, Masters SL, Stirzaker RA, Gorman DJ, Grant RW, Dragoljevic D, Hong ES, Abdel-Latif A, Smyth SS, Choi SH, Korner J, Bornfeldt KE, Fisher EA, Dixit VD, Tall AR, Goldberg IJ, Murphy AJ. 2014. Adipose tissue macrophages promote myelopoiesis and monocytosis in obesity. Cell Metab 19:821–35.

22. Abebe F, Bjune G. 2009. The protective role of antibody responses during Mycobacterium tuberculosis infection. Clin Exp Immunol 157:235–43.

23. Breda J, Banerjee A, Jayachandran R, Pieters J, Zavolan M. 2022. A novel approach to single-cell analysis reveals intrinsic differences in immune marker expression in unstimulated BALB/c and C57BL/6 macrophages. FEBS Lett 596:2630–2643.

24. Watkiss ER, Shrivastava P, Arsic N, Gomis S, van Drunen Littel-van den Hurk S. 2013. Innate and adaptive immune response to pneumonia virus of mice in a resistant and a susceptible mouse strain. Viruses 5:295–320.

25. Chavez JS, Rabe JL, Nino KE, Wells HH, Gessner RL, Mills TS, Hernandez G, Pietras EM. 2023. PU.1 is required to restrain myelopoiesis during chronic inflammatory stress. Front Cell Dev Biol 11:1204160.

26. Patel D, Gaikwad S, Challagundla N, Nivsarkar M, Agrawal-Rajput R. 2018. Spleen tyrosine kinase inhibition ameliorates airway inflammation through modulation of NLRP3 inflammosome and Th17/Treg axis. Int Immunopharmacol 54:375–384.

27. Yadav S, Dalai P, Gowda S, Nivsarkar M, Agrawal-Rajput R. 2023. Azithromycin alters Colony Stimulating Factor-1R (CSF-1R) expression and functional output of murine bone marrow-derived macrophages: A novel report. Int Immunopharmacol 123:110688.

28. Naveen CR, Gaikwad S, Agrawal-Rajput R. 2016. Berberine induces neuronal differentiation through inhibition of cancer stemness and epithelial-mesenchymal transition in neuroblastoma cells. Phytomedicine 23:736–44.

